# Nestin in immature embryonic neurons regulates axon growth cone morphology and Semaphorin3a sensitivity

**DOI:** 10.1101/228296

**Authors:** C.J. Bott, C. G. Johnson, C.C. Yap, N.D. Dwyer, K.A. Litwa, B. Winckler

**Affiliations:** Department of Cell Biology, University of Virginia, Charlottesville, VA 22908, USA.; Department of Anatomy and Cell Biology, The Brody School of Medicine, East Carolina University, Greenville, NC 27834, USA.

## Abstract

Correct wiring in the neocortex requires that responses to an individual guidance cue vary among neurons in the same location, and within the same neuron over time. Nestin is an atypical intermediate filament expressed highly in neural progenitors and is thus used widely as a progenitor marker. Here we show a subpopulation of embryonic cortical neurons which transiently express nestin in their axons. Nestin expression is thus not restricted to neural progenitors but persists at lower levels in some newborn neurons for 2-3 days. We found that nestin-expressing neurons have smaller growth cones, suggesting that nestin affects cytoskeletal dynamics. Nestin, unlike other intermediate filament subtypes, regulates cdk5 kinase. Cdk5 activity is induced by the repulsive guidance cue Sema3a leading to growth cone collapse *in vitro*. Therefore, we tested whether nestin-expressing neurons showed altered responses to Sema3a. We find that nestin-expressing newborn neurons are more sensitive to Sema3a in a roscovitine-sensitive manner, whereas nestin knockdown results in lowered sensitivity to Sema3a. We propose that nestin functions in immature neurons to modulate cdk5 and thereby the Sema3a response. Thus, the transient expression of nestin could allow for temporal modulation of a neuron's response to Sema3a particularly during early axon guidance decisions.

## Introduction

Proper wiring of the nervous system requires that axonal growth cones respond to a variety of extracellular guidance cues to find their correct targets (Kolodkin & Tessier-Lavigne 2011). Semaphorin 3a (Sema3a) is one of many diffusible developmental cues and has been shown to repel axons of responsive neuronal populations (Sibbe et al. 2007). Addition of Sema3a to cortical neuron cultures causes rapid filopodial retraction (within 5 minutes) and ultimately collapse of many axonal growth cones (Dent, E W. 2004). Binding of Sema3a to its receptor neuropilin1 and co-receptor PlexinA leads to downstream activation of the neuronal serine/threonine kinase cyclin dependent kinase 5 (cdk5) (Sasaki et al. 2002; Perlini et al. 2015; Chen et al. 2008). Cdk5 is critical for early neuronal development as a regulator of cytoskeleton, trafficking, and membrane dynamics and, thus, cell morphology (Kawauchi 2014). Sema3a-activated cdk5 phosphorylates multiple substrates to affect the assembly and dynamics of both the actin and microtubule cytoskeleton, and of adhesion components which results in filopodial retraction and growth cone collapse *in vitro* (Sasaki et al. 2002; Ng et al. 2013). Not all cortical neurons are repelled by Sema3a, and the molecular mechanisms for this differential responsiveness are under active investigation (Carcea et al. 2010; Mintz et al. 2009; Ip et al. 2011).

Intermediate filaments are generally thought to provide structure and stability to a cell. In adult neurons, neuronal intermediate filaments, the neurofilaments, play such a structural role to maintain axon caliber (Lariviere & Julien 2003). In addition to the classic neurofilaments, other intermediate filament subunits are expressed during neural development. Vimentin and nestin are both highly expressed in neural progenitor cells (NPC), so much so that nestin expression is widely used as a marker for NPCs. α-internexin and neurofilament proteins expression, on the other hand, is upregulated only after neurons differentiate. Neurofilaments, α-internexin, and vimentin (all expressed in CNS neurons) have also been shown to play additional but poorly understood roles in early neuritogenesis (Shea et al. 1993; Shea & Beermann 1999; Lee & Shea 2014; Walker et al. 2001). The function of nestin in neurons has not been investigated since nestin is not thought to be expressed in neurons.

Little is known about nestin’s function, even in NPCs, where it is highly expressed (Sahlgren et al. 2006), but it appears to offer some protection from cell stresses. However, nestin is often expressed in cancer cells, and has well documented effects on cancer cell migration and invasion by influencing cytoskeletal dynamics and kinase activity (Zhao et al. 2014; Liang et al. 2018; Yan et al. 2016; Hyder et al. 2014). Nestin’s developmental function is best understood in muscle where it acts as a scaffold and regulator for cdk5 at the neuromuscular junction (NMJ). At the NMJ, nestin deletion phenocopies cdk5 deletion (Lin et al. 2005), and nestin is required for agonist induced cdk5 activity (Yang et al. 2011; Pallari et al. 2011; Mohseni et al. 2011). The binding site for the cdk5 complex has been mapped to the unique, long C-terminal tail of nestin (Sahlgren et al. 2003; Sahlgren et al. 2006) which is much larger than conventional intermediate filaments (57 kD vimentin vs ~280kD nestin).

In this work, we show for the first time that nestin has a function in postmitotic newborn cortical neurons. We first characterized the patterns of endogenous nestin expression. As expected, we find that nestin is highly expressed in NPCs. Unexpectedly, nestin protein can be detected during early, but critical, periods of axon formation and is subsequently lost. Secondly, we found that depletion of nestin affects growth cone morphology of immature cortical neurons. Lastly, we discovered that nestin depletion from cultured neurons decreases their sensitivity to Sema3a in a roscovitine-dependent manner. Based on our data, we propose that nestin acts as a “gain control” modulator of Sema3a growth cone signaling during early neuronal differentiation. Since neurons can encounter the same cue at different developmental stages, but respond differently (Kolodkin & Tessier-Lavigne 2011), nestin’s modulation of Sema3a responsiveness might be one mechanism to temporally regulate differential axon guidance decisions in the cortex.

## Results

### Nestin protein expression persists in immature primary cortical neurons in culture

Nestin protein is expressed at high levels in cortical radial glia/neural progenitor cells (NPCs) (Lendahl et al. 1990). In contrast, doublecortin (DCX) and βIII-tubulin expression is low in NPCs but upregulated in postmitotic neurons (Francis et al. 1999). Nestin and DCX/βIII-tubulin are thus routinely used as markers to distinguish these mutually exclusive cell types. E16 mouse dissociated cortical neurons were cultured for 1 day *in vitro* (DIV) and immunostained against nestin and DCX. To our surprise, many DCX-positive cells were also positive for nestin (Figure 1A). These DCX/nestin-double positive cells had the morphology of stage 3 immature neurons (using Banker staging, see methods (Dotti et al. 1988)) (Figure 1A) and were positive for neuronal-specific βIII-tubulin (see Figure 2D), indicating that they had differentiated into neurons. In addition, GFAP (a marker of astrocytes) was not detected at this early stage of development (not shown), in agreement with studies that show gliogenesis, i.e. formation of astrocytes or oligodendrocytes, does not begin until E17/18 or postnatally, respectively (Miller & Gauthier 2007). We note that cells lacking neurites were generally nestin-positive and likely correspond to NPCs and some early stage 1 neurons (see Figure 1D). Cells with short processes which might correspond to stage 2 neurons or to non-neuronal cell types had variable nestin expression.

**Figure 1:**
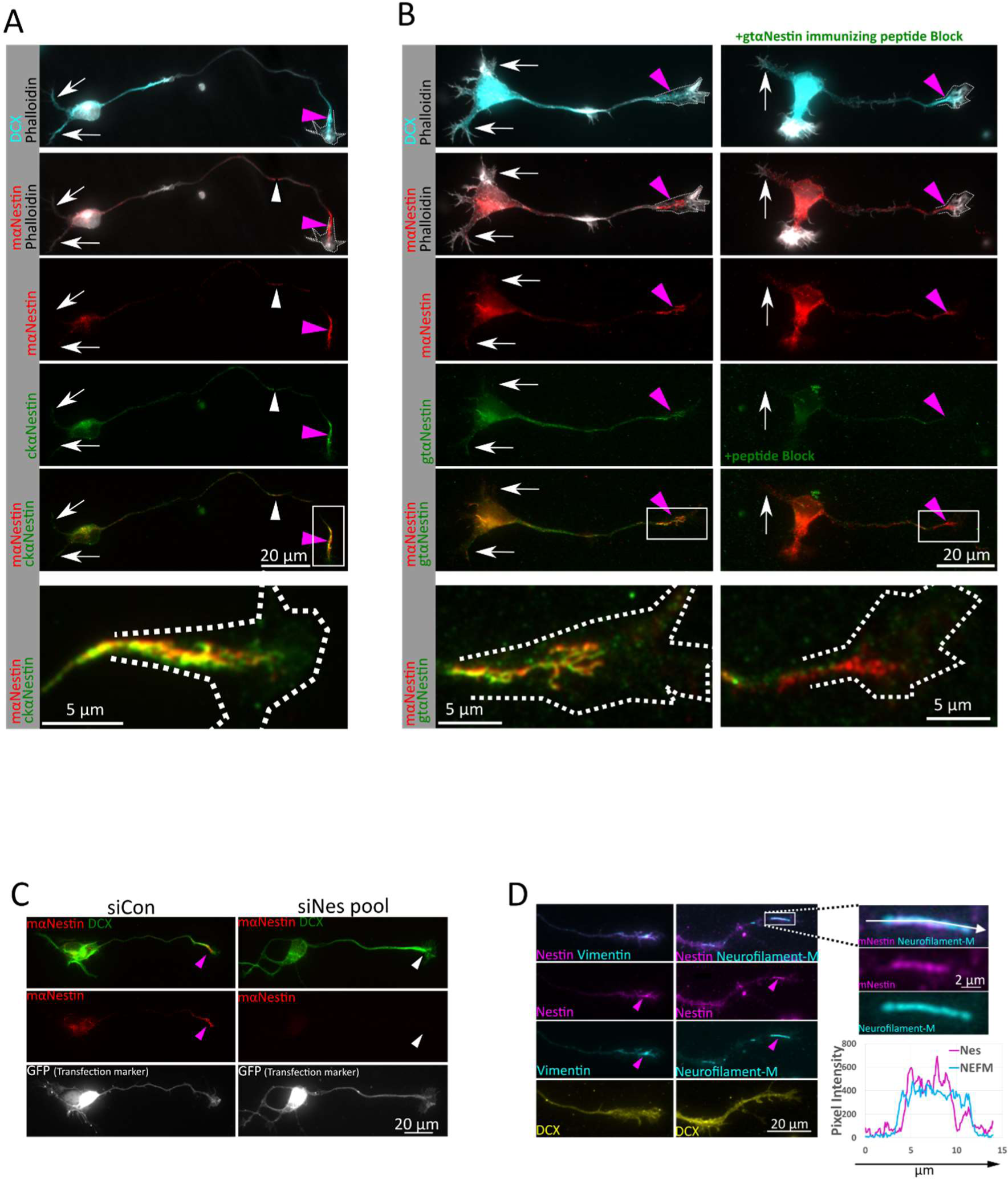
Nestin protein expression persists in immature primary cortical neurons in culture. A. Nestin expression in cultures derived from E16 mouse cortex grown for one day (1DIV). Doublecortin (DCX) immunostaining and a stage 3 morphology was used to identify neurons. Nestin is enriched near the axonal growth cone (pink arrowheads), but largely absent from growth cones of minor dendrites (white arrows) using both mouse and chicken anti-nestin antibodies. White arrowheads indicate short nestin IF segments found along the axon shaft. Bottom panel is enlarged inset of growth cone, rotated, from within the white box. Growth cone outline drown from the phalloidin stain shown in the panels with phalloidin and in the enlargement. B. Validation of nestin antibody staining on stage 3 neurons with mouse and goat anti-nestin antibodies with immunizing peptide antibody blocking. Both antibodies again show a distal axon enrichment of nestin (pink arrowheads) while the dendrites again are not labeled (white arrows). Pre-incubation of the goat anti-nestin antibody with the immunizing peptide abolishes all staining but staining in the distal axons is still detectable with the mouse anti-nestin antibody which was raised against a different epitope. Shown are E16 mouse cortical neurons 1DIV. Bottom panel is enlarged inset of growth cone from within the white box. Growth cone outline drown from the phalloidin stain shown in the panels with phalloidin and in the enlargement. C. Nestin siRNA results in loss of immunostaining, validating that the axonal growth cone staining is due to nestin protein. GFP was co-transfected as a transfection indicator. After 36 hours in culture, cells were fixed and immunostained for nestin and DCX. The number of GFP positive cells positive for nestin in the axon as a fraction of the number of cells counted was quantified. D. Nestin is co-localized with other intermediate filaments in the same assembly group (vimentin and Neurofilament-Medium NF-M) in the distal axon (pink arrowheads) of E16 mouse cortical neurons 1DIV. N-FM expression was low in immature cortical neurons cultured for only 1DIV. High magnification of the boxed region and linescans reveal that nestin and N-FM appear to decorate the same filament structure in partially co-localizing subdomains, as expected for nestin containing intermediate filaments. Pink arrowheads indicate nestin positive axonal intermediate filament heteropolymers.

**Figure 2:**
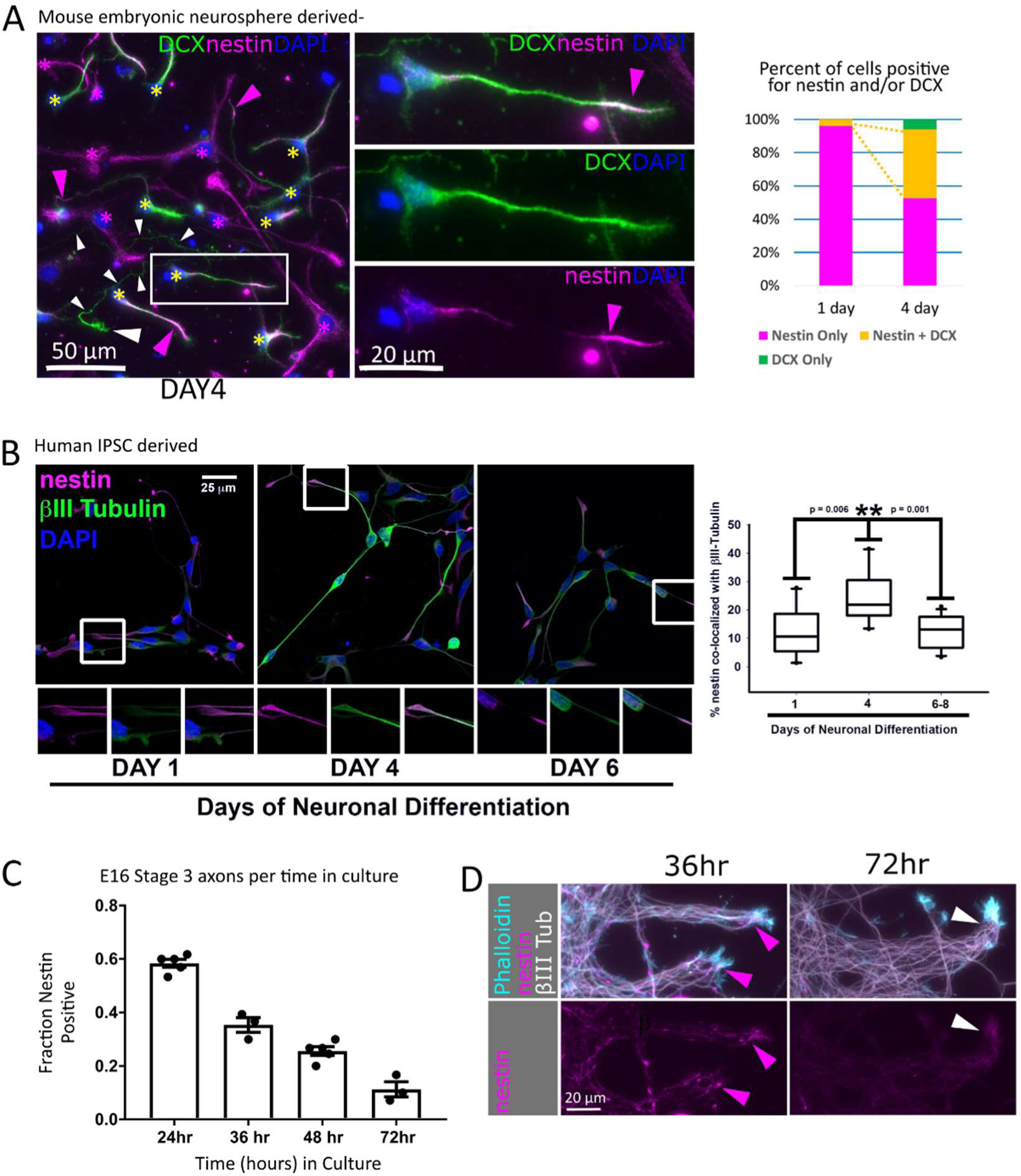
Nestin-expressing neurons are observed in multiple rodent and human culture models. A. Mouse Neurosphere NPC cultures: Nestin expression was assessed in differentiating dissociated neural progenitor cells (mouse NPC) after 4 days of neuronal differentiation. DCX-positive neurons frequently stain positive for nestin (yellow asterisk), especially in axon tips (pink arrowheads). An example of axon positive for DCX but not nestin is marked with a series of white arrowheads along the length of the long and thin axon, and a large white arrowhead indicating the growth cone. Nestin single positive cells are marked with a pink asterisk. One example of a double-positive cell is shown in the middle panels to highlight its neuronal morphology and tip-enriched nestin staining (pink arrowhead). The proportion of cells positive for only nestin (red bar), only DCX (green bar) or double-positive for DCX and nestin (yellow bar) was quantified for NPCs cultured under differentiation conditions from 1 or 4 days. Yellow dotted lines highlight the expanding double positive population (yellow bar). 323 cells were counted at 1 DIV, and 219 cells counted at 4 DIV. B. Human IPSC derived neuronal cultures: Nestin is expressed in human (IPSC) differentiating neurons derived from NPCs in dissociated cultures under differentiating conditions from 1, 4 and 6 days. The distribution of nestin is similar to the mouse NPC-derived neurons described above. Percent co-localization with the neuronal-specific βIII tubulin was quantified. A close-up of a nestin-positive axon tip in the boxed region is shown in the panels below the respective image. n=14 (day 1), 10 (day 4), and 14 (day 6-8). (Statistics: Mann-Whitney t test) C. Mouse primary neurons cortical neuron cultures: Percentage of nestin-positive neurons rapidly decreases with time in culture. (30-60 stage 3 neurons were counted per time point for 3 to 5 experiments as shown as the n). Cortical neuron cultures were prepared from E16 mice embryos and culture times are indicated. D. Mouse primary explants: Axonal nestin expression is progressively lost between 36 to 72hours in culture. Significant axonal nestin immunostaining is no longer detected by 72 hours. E16 mouse cortex was explanted by incomplete dissociation and cultured. Axon fascicles emerging from the explant are shown. Arrowheads point at axon tips.

Because nestin is considered a marker of NPCs, we performed a series of rigorous controls to validate the observed staining. We obtained two additional nestin antibodies (raised in chicken (Chk) or goat (Gt)), all against different epitopes in the C-terminal region of nestin. Both antibodies produced immunostaining patterns similar to the mouse anti-nestin antibody (Figure 1A,B). A blocking peptide was available for the goat anti-nestin antibody which allowed testing the specificity of the observed axonal nestin staining. Pre-incubation of the goat anti-nestin antibody with the immunizing peptide prior to labeling specifically reduced immunostaining with the goat antibody, but nestin was still detected in distal axons with the mouse anti-nestin antibody in the same cell (Figure 1B). Both of these antibodies appear highly specific to nestin since they detected single bands (~300kd) by immunoblot (consistent with the expected size of nestin) in DIV1 E16 cortical mouse neuron lysate (Fig S1B).

As an additional validation of antibody specificity, we performed nestin knockdowns with either control, non-targeting siRNA (siCon) or a nestin-specific pool of 4 siRNAs (siNes-pool) using Amaxa nucleofection prior to plating dissociated neurons. GFP was co-electroporated to visualize transfected cells (Figure 1C). Cultures were fixed after 36 hrs and stained for nestin. The intensity of axonal nestin was then quantified. At 36 hrs, approximately 1/3 of GFP-positive stage 3 neurons stained positive for nestin after siCon electroporation, similar to untransfected cells (see below Figure 2C), whereas nestin staining was significantly reduced and essentially undetectable in GFP-positive cells electroporated with siNes-pool (Figure 1C). The specificity of the three nestin antibodies was also assessed by western blot after siRNA knockdown in neurons (Figure S1A). Since electroporation is not very efficient in neurons after only 36 hours, the knockdowns were not complete, but decreased nestin levels were found after siNes-pool nucleofection for all three antibodies confirming that they were specific for nestin. When mouse nestin-myc was transfected into HEK293 cells together with either siCon or siNes-pool siRNA, a near complete loss of nestin signal was observed in the siNes-pool treated condition, while the related IF vimentin was not affected (Figure S1A’). We conclude that nestin protein is indeed expressed in immature neurons in culture. Close reading of the literature revealed that nestin protein has been detected in neurons in human and mouse, both in early differentiating neurons (Walker et al. 2007; Yan et al. 2001; Cattaneo & McKay 1990; Crews et al. 2011; Messam et al. 2002; Messam et al. 1999), and in subpopulations of adult neurons (Gu et al. 2002; Hendrickson et al. 2011; Decimo et al. 2011; Guo et al. 2014; Liu et al. 2018). Recent advances in single cell western blotting have revealed persistent nestin expression in βIII tubulin positive differentiated neural stem cells (Hughes et al. 2014).

Despite the evidence for nestin protein expression not being exclusive to neural stem cells, nestin continues to be used as a specific neural stem cell marker. We thus performed a side by side comparison (Figure S1C) of nestin staining in cultures counterstained with the NPC marker Sox2 and the neuronal marker DCX to compare the relative levels of nestin protein. This comparison within a single imaging field demonstrates that the nestin protein immunofluorescence level seen in axons (high DCX, but low Sox2 expression) was many times lower than nestin expression in an NPC cell body (high Sox2, but low DCX expression). High levels of anti-nestin antibody staining was thus confirmed to be a reliable NPC marker, when used as a tool to label NPCs and is clearly distinguishable from the low axonal levels of nestin.

### Nestin is present in filaments in distal axons and growth cones

Having validated the nestin antibodies, we next characterized endogenous neuronal nestin distribution. Nestin was not uniformly distributed, but primarily localized to the distal region of the growing axon (pink arrowheads) and was rarely observed in secondary neurites (white arrows) (Figure 1A). Variable levels of nestin were often also detected in the cell body. In addition to the distal axon accumulation of nestin, short linear nestin profiles (called squiggles) were often found distributed along the length of the axon (white arrowhead) (Figure 1A), which is typical of neurofilaments in young neurons (Benson, D. L., et al. 1996; Wang et al. 2000). Nestin is an obligate heteropolymer and requires another cytosolic IF type to assemble into filaments (Steinert et al. 1999). If nestin existed as assembled filaments in neurons, it should be part of a filament composed of other IF subunits in addition to nestin. Indeed, we found that nestin colocalized with vimentin, neurofilament-medium (NF-M) (Figure 1D), α-internexin and neurofilament-light (NF-L) (not shown). Neurofilament heavy (NF-H) was not detected at these early time points, as NF-H is expressed in more mature neurons (Benson, D. L., et al. 1996). A magnification inset and a linescan show that nestin and NF-M occupy subdomains within a heteropolymer, as has been demonstrated with nestin filaments in other cell types (Leduc & Manneville 2017).

In addition to assembled nestin-containing filaments (squiggles) along the axon, we also see nestin-positive staining in the growth cone central domain which extends to various degrees more peripherally. A typical pattern is represented by the growth cone shown enlarged at the bottom of Figure 1A. Nestin staining appears consolidated at this magnification and resolution. Individual filaments cannot easily be discerned. This staining may represent a tight bundle or tangle of short nestin-containing intermediate filaments (squiggles). In more spread growth cones (such as the growth cone shown enlarged in the bottom left of Figure1B) nestin staining appears clearly filamentous, suggesting that nestin in neurons can exist at least partially as an assembled heteropolymer filament.

### Nestin-expressing neurons are observed in multiple rodent and human culture models

Since neurons are not usually thought to express nestin, we looked at other neuronal models to see if we observed DCX/nestin double positive neurons in other contexts. When we prepared neuronal cultures from rat cortex, we also observed DCX/nestin double positive neurons (not shown). Next, we grew mouse NPCs as neurospheres and then differentiated them into neurons in culture in order to determine whether neurons differentiated from NPCs continued to express nestin. Differentiation into neurons takes several days using this protocol. One day after neurosphere dissociation and plating, >95% of cells were nestin-positive, but did not express DCX, consistent with them being NPCs (Figure 2A). The remainder had started to differentiate into neurons and expressed both DCX and nestin. After 4 days of differentiation in dissociated culture, ~50% of cells had differentiated into neurons (DCX-positive). A large proportion of the DCX-positive neurons also expressed nestin. The DCX/nestin double positive cells had a morphology similar to stage 2 or stage 3 cultured primary neurons and cells with a stage 3 morphology had nestin localized to the tips of neurites (Figure 2A inset). Only around 10% of cells had no detectable nestin immunopositivity (Figure 2A) and expressed only DCX, consistent with being more differentiated neurons. Differentiating NPCs after 4 days became too crowded to assess on a cell-by-cell basis.

To test whether human neurons similarly express nestin during early differentiation, we assessed nestin expression in cultures derived from human neural progenitor cells derived from induced pluripotent stem cells (IPSCs). At 1 day, nestin expression is high and βIII tubulin expression is low, resulting in low co-localization of nestin and βIII tubulin. After 4 days of differentiation, nestin colocalization with βIII tubulin peaked, and nestin staining was clearly detected at the tips of βIII-tubulin positive axons (Figure 2B), confirming nestin expression also in cultures of human neurons. Co-localization again drops at ~ 1 week due to decreasing nestin expression, however nestin is still expressed at the tips of some neurites. To assess if nestin expression occurred in neurons that are differentiating in a more tissue-like context, we also grew human NPCs as large cortical spheroids, often referred to as “minibrains” (Figure S2A). Minibrains recapitulate several developmental processes which are also observed during *in vivo* development. Nestin staining was clearly detected in neurons (identified by βIII-tubulin expression). Nestin staining was most prominent in neuronal processes with lowerβIII-tubulin expression (less mature neurons), while cells expressing higher levels of βIII-tubulin (more mature neurons) were mostly negative for nestin. We conclude that nestin is transiently expressed in neurons in both human and rodent neurons, and can be observed in a model that recapitulates human cortical development.

### Nestin protein is transiently expressed broadly among cortical neuron cell types but decreases with maturation

Typical techniques to culture primary neurons select for cells that just recently completed mitosis, or are undergoing migration at the time of dissociation (Banker & Cowan 1977). We thus wondered if nestin-positive neurons represent a transition state between nestin-positive neural progenitor cells (NPCs) and nestin-negative neurons. Alternatively, nestin-positive neurons could be a stable neuronal cell type which maintain nestin expression over longer times in culture. Since cultured cortical neurons maintain cell type fates after dissociation (Romito-DiGiacomo et al. 2007; Digilio et al. 2015), we used two cell-type specific transcription factors to distinguish cortical-projecting (Satb2-positive) from subcortical-projecting (Ctip2-positive) neurons. Nestin expression in DCX-positive neurons did not correlate with either Ctip2 or Satb2 expression (Figure S2B). Similarly, when cultures were prepared from E14, E15 or E16 embryos (representing different cortical layer fates), about 2/3 of DCX-positive neurons were nestin-positive after 24 hours in culture (data not shown), regardless of embryonic age at dissection. These data suggest that nestin is expressed by immature neurons of all cortical layers.

Since nestin-positivity was not skewed by neuronal cell type, we next assessed if nestin expression was correlated with time after differentiation. To this end, we utilized an *in vitro* time course of neurons dissociated from E16 mouse cortex (Figure 2C), since time in culture represents the maturity of the cell. For the first 12-24 hours, around two-thirds of stage 3 neurons expressed nestin in the axon. Plating density had no effect on the percentage of neurons that express nestin (not shown). Nestin expression dropped steadily after 24 hours, with only one third remaining nestin-positive at 36 hours, one quarter by 48 hours, and less than a tenth by 72 hours (Figure 2C). The change in axonal nestin staining is easily appreciated in axon bundles growing from a cortical explant. At 36 hours, nestin expression was clearly seen mostly in the distal region of some axons, but was reduced to near background levels at 72 hours (Figure 2D). These data demonstrate that nestin is expressed transiently in a substantial subpopulation of differentiating cortical axons, and subsequently downregulated as differentiation proceeds.

### Nestin is expressed in subpopulations of developing cortical neurons in vivo

We next sought to determine if there was an *in vivo* correlate to the axonal nestin expression we observed in cultured neurons. Others have shown that developing cortical neurons in the intermediate zone (IZ) consist of a mixture of axons of variable states of maturation-pre-existing axon tracts laid down by earlier pioneer neurons, and later born neurons which initiate axon projections during migration through the IZ (Namba et al. 2014). We thus imaged axons in the lower intermediate zone (IZ) of E16 mouse developing cortex, similarly to where nestin mRNA was detected by Dahlstrand (1995). *In vitro*, nestin was not present in all axons and did not fill the whole length of an individual axon. We thus predicted that nestin positive axons would be detected as a subpopulation of axons in the IZ. We also predicted that axonal nestin would be lower than nestin in NPCs/radial glia.

For orientation, a low magnification overview of one hemisphere of the cortex showing vimentin and α-internexin (INA) expression is shown in Figure 3A, along with a schematic. The boxed region in Figure 3A indicate the lateral lower IZ which is the region in the schematic imaged in the following panels. All panels in Figure 3 B-E have the radial glia oriented vertically and axon tracts oriented horizontally. INA is an intermediate filament expressed early in neuronal development, but not expressed in radial glia (Kaplan et al. 1990; Fliegner et al. 1994; Benson, D. L., Mandell J. W., Shaw G. 1996; Dahlstrand et al. 1995). As expected, INA is readily detected in axons of the IZ (Figure 3A,B). Vimentin is expressed in both radial glia and, at a lower level, neurons (Cochard & Paulin 1984; Boyne et al. 1996; Toth et al. 2008; Yabe et al. 2003).

**Figure 3:**
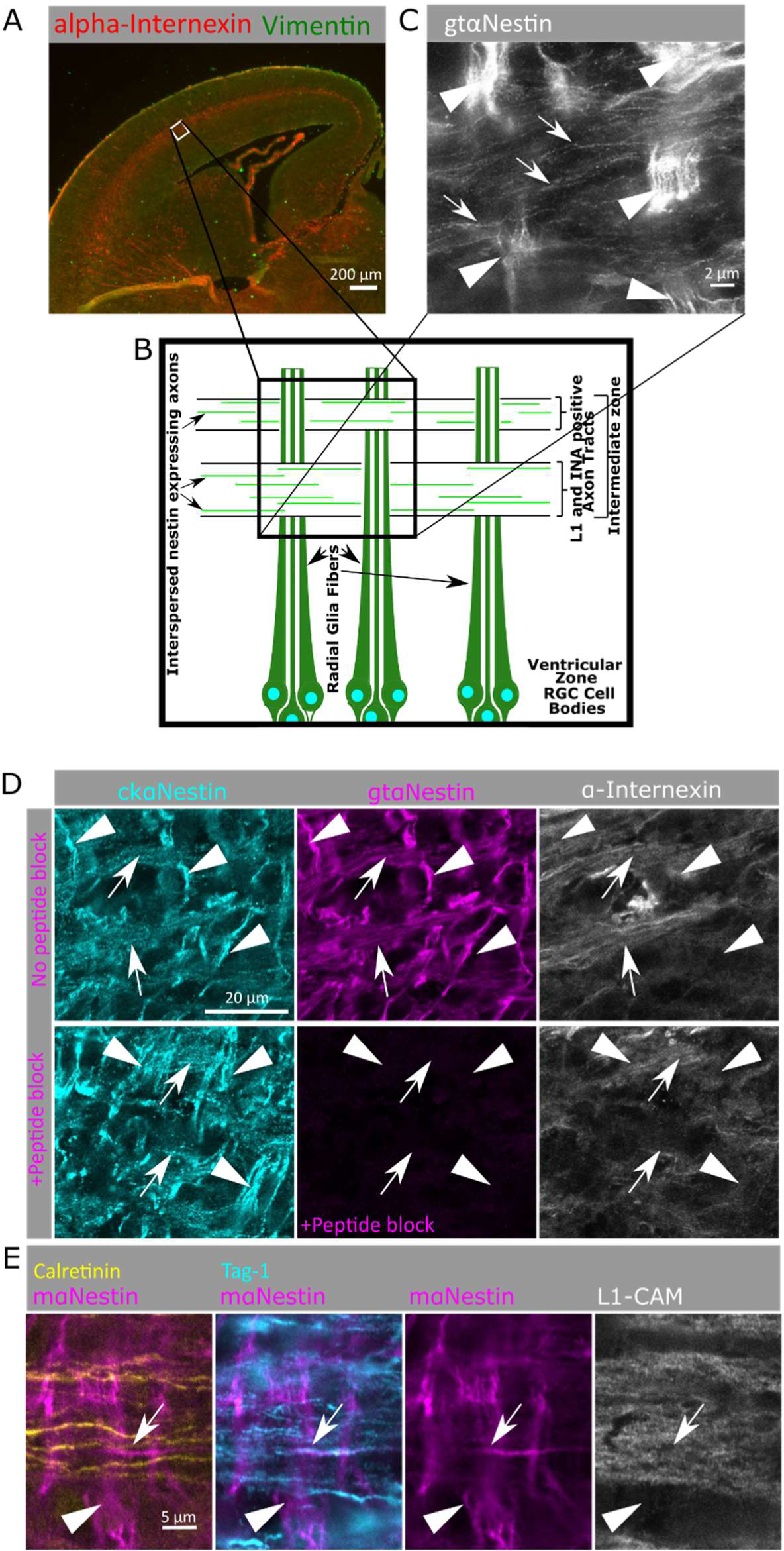
Nestin is expressed in sub-populations of cortical axons in developing cortex. A. A low magnification of a coronal cortical section (E16 mouse) is shown for orientation (top) and showing α-internexin localization to the axon rich intermediate zone. The white box (lateral lower Intermediate Zone) is the region imaged in the following panels, and is (B) diagramed below illustrating the vertically oriented radial glia, and horizontal IZ axon bundles. C. Super resolution microscopy (STED) of nestin immunostaining reveals bright staining of radial glial fibers (arrowheads) as well as fainter staining of axons (arrows) in mouse E16 cortex. Many axons in the axon fascicle do not express nestin, so only a subset of axons in the intermediate zone express nestin at this time point. Arrow heads indicate radial glia and arrows indicate nestin-positive axons. D. Nestin staining of the lower intermediate zone of E16 mouse cortex using chicken anti-nestin (cyan) and goat anti-nestin (magenta) antibodies. Axon tracts are visualized with α-internexin antibody (white). Nestin staining is found in radial glia fibers (arrowheads) as well as in α-internexin-positive axon tracts (arrows). The goat anti-nestin antibody was preincubated with immunizing peptide on sequential cryosections in the lower panels. All staining with the goat anti-nestin antibody was blocked by peptide preincubation, including the axon tract staining, demonstrating that the axon staining was specific and not background staining. All images correspond to higher magnification of the lower intermediate zone of the lateral E16 mouse cortex (boxed regions in A). Radial glia are oriented vertically (arrowheads) and axon tracts are oriented horizontally (arrows). D. Identification of nestin-positive axons in mouse cortex. The L1-CAM-positive axons (white) contain mixed populations of both cortical and thalamic projections in this brain region. Nestin (magenta) is specifically expressed in axons originating from the cortex (Tag-1-positive; cyan), but not in axons with thalamic origin (calretinin-positive, yellow). Arrow heads indicate radial glia and arrows indicate nestin-positive and Tag1-positive axons.

As expected, nestin staining intensity in the radial glia (arrowheads in Figure 3C,D,E) was high, which made evaluation of nestin staining in axon tracts challenging. However high resolution confocal microscopy, the use of distinct axonal markers, and sequential imaging of four channels permitted us to resolve and distinguish between the segregated but interwoven cellular processes of radial glial and IZ axons. Staining with chicken and goat anti-nestin antibodies labeled the bright radial glia fibers (INA negative) oriented vertically (arrowheads) (Figure 3D). Similarly to Dahlstrand’s (1995) findings for nestin mRNA, INA is present in early cortical axon tracts (arrows), and nestin immunoreactivity was clearly detected in these axon tracts, albeit at lower levels compared to radial glia fibers. This staining pattern is similar to what is seen with *in vitro* axon bundles (Figure 2D). Early defining work had observed low nestin staining on spinal cord axons, but attributed it to background staining by the antibody (Hockfield & McKay 1985). To assess the specificity of nestin staining on IZ axons, we repeated the peptide blocking assay used in Figure 1 for the goat anti-nestin antibody on E16 cryosections (Figure 3D). The staining by the goat nestin antibody was completely blocked by pre-incubation with the immunizing peptide, both on radial glia fibers and on axon tracts, indicating specific staining of nestin on axons *in vivo*. Both radial glia fibers and axons were still stained with the chicken nestin antibody. In addition, specificity of both the mouse 29178 antibody and the goat antibody to nestin was assessed by immunoblot of E16 mouse cortex lysate, and resulted in clearly labeled single bands that corresponds to the expected size of the nestin protein (Fig. S1B’).

To further characterize the IF content of the lower IZ axons, co-staining for nestin and vimentin was performed (Figure S3A,B). Vimentin was found to have a similar staining pattern to nestin, with high levels in radial glia, and low levels in axons. The nestin- and vimentin-positive IZ fibers were also positive for the axon marker L1-CAM (Figure S3A,B), confirming their identity as axons. The same axonal nestin staining pattern was seen with the mouse anti-nestin Rat401 monoclonal antibody, the initial antibody used to characterize and define nestin (Hockfield & McKay 1985) (Figure S3B). Thus, multiple pieces of evidence demonstrate that nestin is expressed in axons in the IZ at E16 in mouse cortex.

To better visualize the axonal nestin, we performed STED super-resolution microscopy on goat anti-nestin immunostained E16 mouse cortex. The bright nestin-positive radial glia fibers were again the most prominently stained feature, but individual axons with less bright nestin staining could be unambiguously resolved. Nestin is clearly present on a subset of axons of the IZ (Figure 3C). In addition to the axons of cortical neurons (Tag-1 positive) of variable maturity, the IZ contains axons of neurons projecting from the thalamus, calretinin-positive thalamo-cortical neurons (Takebayashi et al. 2015). We thus used these markers to further define the nestin expressing sub-populations (Figure 3E): L1-CAM for all axons, TAG-1 for cortical axons, and calretinin for thalamocortical axons. Intriguingly, we observed that a subpopulation of cortical (TAG-1 positive) axons had detectable nestin staining whereas the calretinin-positive (thalamo-cortical) axons lacked nestin staining (Figure 3E). However, we note that this could be due to differences in the cell type, and/or maturity level of cortical vs thalamic originating axons.

### The level of nestin expression in neurons influences growth cone morphology and response to Sema3a

An important question is whether nestin plays a functional role in axons, or is simply left over from the mother NPC and no longer plays a role. As a starting point for probing whether nestin still has a function in neurons, we determined if axons and growth cones were the same or different in neurons with (+) and without (-) nestin (representative images in Figure 4A). We measured axon length, growth cone area, and number of growth cone filopodia (diagram in Figure S4F). Nestin levels were not correlated with any variance in axon length (Figure 4E), but interestingly growth cone area was significantly smaller when nestin was present (Figure 4F). This finding is consistent with previous reports in frog neurons, where the neurons expressing the nestin homolog, tannabin, had smaller blunted growth cones (Hemmati-Brivanlou et al. 1992). How growth cone size is regulated and what effect growth cone size has on growth cone functions is unknown, but other molecules that regulate the actin and microtubule cytoskeleton display a similar phenotype. Growth cone size therefore may be representative of the underlying overall cytoskeletal dynamics, where a more stable cytoskeleton leads to larger growth cones, and a more dynamic cytoskeleton leads to smaller growth cones (Khazaei et al. 2014; Poulain & Sobel 2007). Others have noted a correlation between axon growth rates and growth cone size, however our measurements of axon length do not show any significant differences (Argiro A., et al. 1984; Ren & Suter 2016).

**Figure 4:**
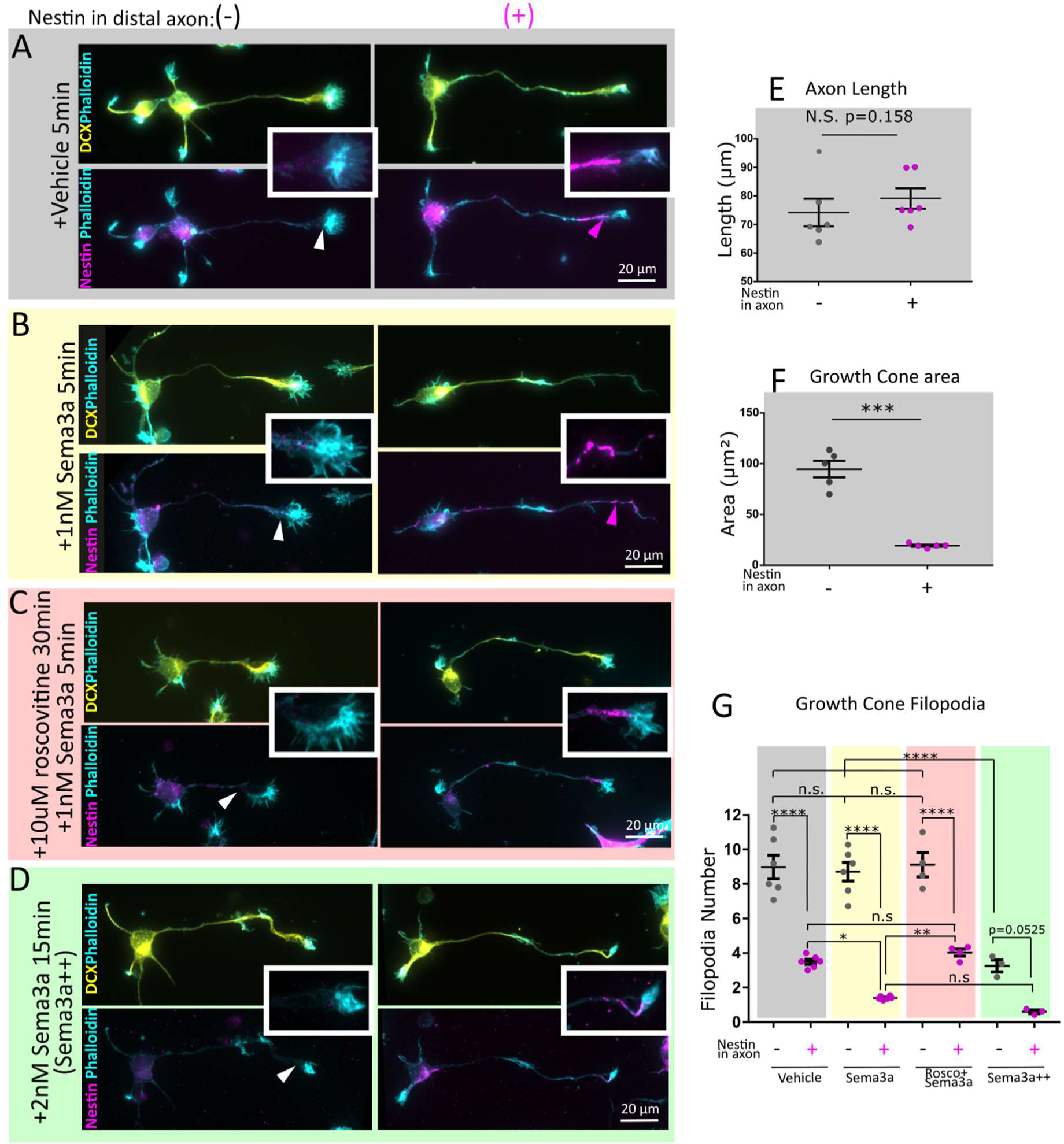
Nestin-expressing neurons have smaller growth cones with fewer filopodia, and are more sensitive to Sema3a in a cdk5-dependent manner. A. E16 mouse cortical neurons 1DIV (stage 3) were immunostained against DCX (to confirm neuronal identity), phalloidin (to visualize the growth cones), and nestin. Cells were characterized based on nestin immunoreactivity in the distal axon nestin as Nes(-)-not detected, Nes(+)-nestin positive. Condition are untreated (A), 1nM sema3a 5 minutes (B), pretreatment with 10µm Roscovitine 30 minutes prior to 1nM sema3a treatment for 5 minutes (C), and a Sema++ treatment of 2nM Sema3a for 15 minutes (D). Insets allow for appreciation of the filamentous nature of the nestin immunostaining and proximity to the growth cone. Channel intensity is adjusted equally for nestin and DCX. In the low magnification, phalloidin labeling has been adjusted variably to show morphology, but in the insets the phalloidin channel was adjusted in an equal manner. E,F. Nestin expressing neurons are not significantly different by axon length (E), but have significantly smaller axon growth cones (F). Neuron morphology was quantified (30-40 cells per experiment for 6 (E), or 5 (F) independent experiments). Mean of each experiment and SEM are indicated. Statistical test was t-test with n=6 (E), or n=5 (F). G. Neurons with nestin have fewer filopodia compared to those without nestin. In cells treated with 1nM Sema3a for 5 minutes, the nestin positive cells had a significant reduction in filopodia compared to untreated nestin positive cells, whereas the nestin negative cells did not. Conditions have been color-coded for ease of comparison. Roscovitine inhibited the decrease in filopodia number after Sema3a treatment. When a high dose (2nM) and longer treatment (15minutes) of Sema3a is used, both nestin negative and positive neurons retract filopodia and significantly so, indicating nestin is not absolutely required for Sema3a induced filopodial retraction, but increases Sema3a sensitivity. Filopodia number were quantified for 6 (vehicle, Sema3a), 4 (roscovitine + Sema3a), or 3 (Sema++) independent experiments. The average of 30-40 cells quantified per experiment is shown. Mean of each experiment and SEM are indicated. Normality was confirmed with the Shapiro-Wilk normality test. Statistical test was one way ANOVA with Tukey’s multiple comparison correction, with n= number of experiments. Each condition was compared to every other condition.

Since nestin regulates cdk5-dependent processes in muscle cells, we wondered if nestin could affect cdk5-dependent processes in neurons as well. Sema3a is upstream of cdk5 in neurons (Kobayashi et al. 2014), and we thus utilized an *in vitro* bath application “Sema3a collapse assay” to test this idea. In cultured neurons, phosphorylation of cdk5 substrates and morphological changes in growth cones (Perlini et al. 2015; Dent 2004) start within minutes of Sema3a exposure with the loss of growth cone filopodia, resulting ultimately in a collapsed growth cones with no filopodia.

However, prior to complete growth cone collapse, Sema3a leads to retraction of individual filopodia (Dent 2004). Given nestin’s role in promoting cdk5 activity at the neuromuscular junction, we hypothesized that growth cone filopodial retraction in response to Sema3a occurred in a nestin dependent manner. In order to quantitatively address this, we counted actin-rich filopodial protrusions of the growth cones (Figure 4G). In untreated cells, filopodia number was inversely correlated to axonal nestin levels (Figure 4G). After a low dose of Sema3a treatment (1nM for 5 minutes) (Perlini et al. 2015), nestin-positive cells retracted growth cone filopodia significantly, whereas nestin-negative growth cones were not significantly affected by Sema3a during the 5 minute timeframe (shown in Figure 4B, quantified in Figure 4G). Area of the growth cone was not significantly altered in this short time frame as expected (Dent 2004), so we relied on the retraction of filopodia as our most sensitive measurement of Sema3a sensitivity. Because Sema3a is known to act in part through cdk5, we added 10µM of the cdk inhibitor roscovitine for 30 minutes prior to Sema3a application. Roscovitine pretreatment inhibited the decrease in filopodia number after Sema3a treatment (Figure 4C,G). Roscovitine inhibits cdk5 potently, but it also inhibits some of the mitotic cdks (cdk2 and cdc2), albeit less effectively. Since neurons are post-mitotic, they have little to none of the mitotic cdk activities. It is thus likely that the roscovitine inhibition of the Sema3a effect is due to cdk5 inhibition.

Lastly, nestin is not absolutely required for Sema3a sensitivity since higher and longer Sema3a stimulation (2nM for 15 minutes) resulted in filopodial retraction even in nestin negative cells (Figure 4D,G). Together these data demonstrate a strong correlation between axonal nestin expression and growth cone size, and Sema3a mediated filopodial retraction.

### Nestin depletion results in abnormal growth cone morphology and resistance to Sema3a

To test for nestin dependent sensitivity to Sema3a directly, we used nestin siRNAs to deplete nestin protein from neurons and assess growth cone morphology and response to Sema3a. Neurons electroporated with the siNes-pool and GFP (to visualize transfected cells) (Figure 1C, S1A, Figure 5A,E) were analyzed for differences in morphology. Neurons that received the siNes-pool had no variance in axon length (Figure 5C), as expected, but had large growth cones (Figure 5D) and more growth cone filopodia (Figure 5F) compared to siCon, mirroring the effects seen from analysis of endogenous variation of nestin expression (Figure 4). To confirm that this effect was specific to nestin depletion and not an off-target effect, the 4 siRNAs that make up the siNes-pool were tested individually. Two of them (#1 and #17) were effective for both depleting nestin as well as for increasing growth cone size and filopodial number (Figure S4A-E). More than one distinct siNes thus resulted in the same phenotype as the siNes-pool, consistent with specificity of the phenotype to nestin depletion. Consistent with this conclusion, siNes #4 failed to reduce nestin levels and did not affect growth cone morphology, similar to siCon (Figure S4A-E).

**Figure 5:**
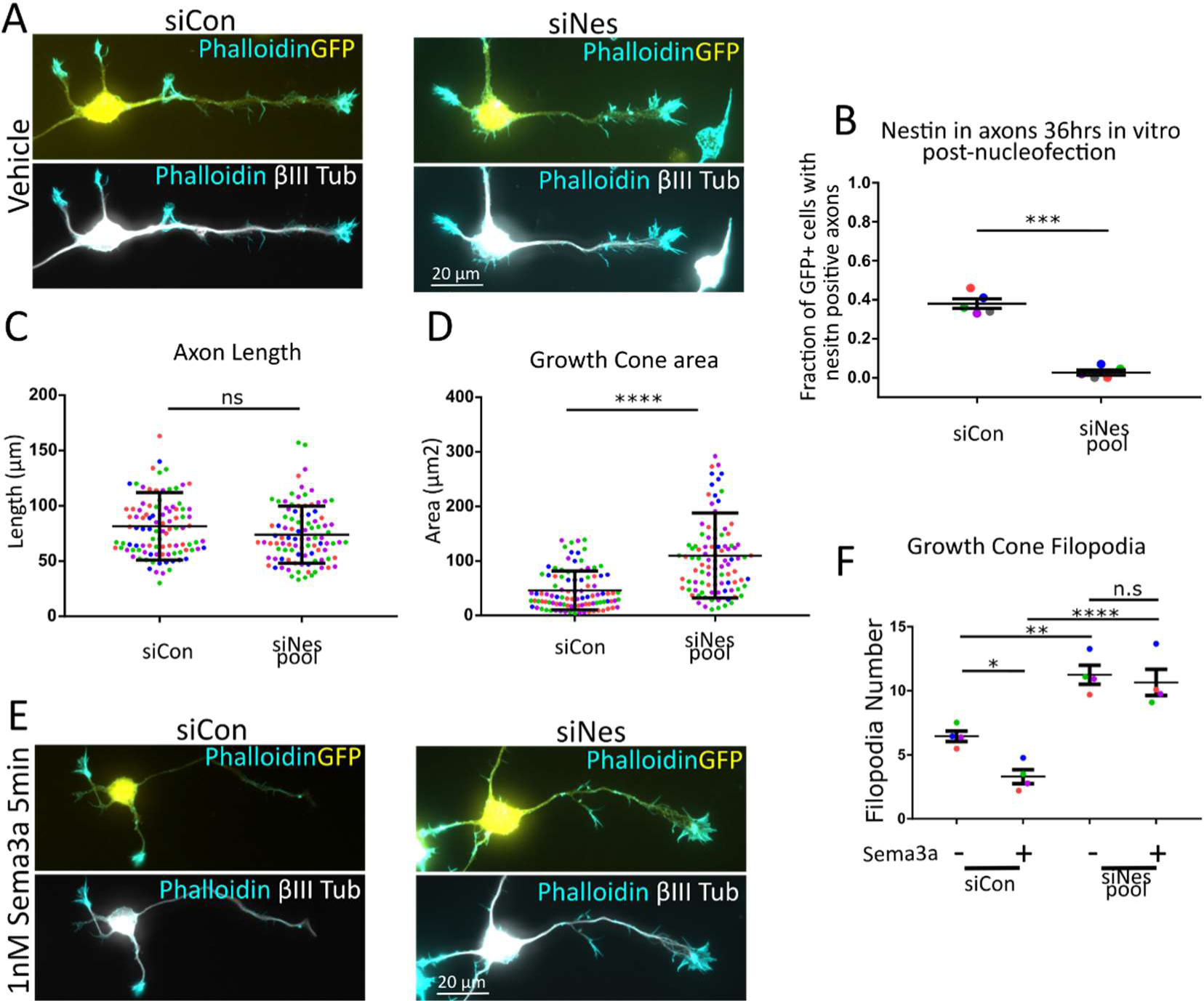
Nestin siRNA results in abnormal growth cone morphology and resistance to Sema3a. Representative images of primary E16 Mouse cortical neurons that were co-transfected with a plasmid encoding GFP and either control siRNA (siCon) or nestin siRNA (siNes) and cultured for 36hrs. GFP booster was used to visualize transfected cells, Tuj1 (βIII-tubulin) was used to confirm neuronal identity and for cell morphology, and phalloidin to visualize growth cones (A,E). The efficiency of nestin knockdown was quantified (B) by counting GFP (transfected) and nestin positive cells by immunostaining was quantified in stage 3 neurons. ~1/3 of siCon neurons express nestin at 36 hrs in the axon. Very few transfected neurons showed nestin expression in the axon under siNes conditions. 120-140 cells per condition were counted in n=5 independent experiments. p< 0.001 (Paired t-test). In vehicle treated conditions (A), siNes treated cells had no significant difference in axon length (C) but resulted in significantly larger growth cones (D). n= ~100 cells (plotted on the graph) were measured for each condition and significance measured by unpaired Mann-Whitney rank comparison test. Shapiro-Wilk normality test indicated the data were not normal. A t-test was also performed (to be consistent with other analysis) for the growth cone area data separately with each experimental average, n=4 (each a different color on graphs), and resulted in a significant p value of 0.025. siCon or siNes transfected neurons were treated for 5 minutes with 1 nM Sema3a (E). siNes resulted in more filopodia per growth cone, and these filopodia were resistant to Sema3a retraction (F). ~100 or cells in 4 independent experiments were averaged and n=4 was used for statistical analysis (one way ANOVA with Sidak correction – only the pairs shown were compared). Normality was confirmed with the Shapiro-Wilk normality test.

As expected, siCon transfected neurons had a significant reduction in filopodia number following 1nM Sema3a treatment for 5 minutes. Neurons transfected with the siNes pool and then treated with Sema3a (1nM 5 minutes), in contrast, showed no change in filopodia number (Figure 5E) and they were thus insensitive to this dose of Sema3a. These results demonstrate, surprisingly, that nestin greatly increases the sensitivity of axonal growth cones to Sema3a.

## Discussion

Neural development requires that axonal growth cones respond to a variety of extracellular guidance cues to change their direction of growth for proper wiring of the nervous system (Kolodkin & Tessier-Lavigne 2011). Importantly, not all growth cones traversing the same environment respond the same way to the same cues at the same time (Carcea et al. 2010; Mintz et al. 2009). Even more remarkably, the same growth cone often responds differently to the same axon guidance cue when encountering it at a different point in time and space (Kolodkin & Tessier-Lavigne 2011). As an axon grows into new brain regions, it must therefore not only respond to new cues, but often differently to the same, previously encountered cue. The molecular mechanisms underlying this variable response to the same cue are poorly understood. We propose that nestin acts as a sensitizer and might provide a gain control mechanism, such that axons lacking nestin are less repelled by Sema3a than axons expressing nestin.

### Nestin is highly expressed in NPCs but expression persists in young neurons

We made the surprising discovery that nestin, the most commonly used marker for NPCs, is also found in young neurons. Our conclusion that nestin is in fact expressed in neurons is based on multiple lines of evidence, including multiple antibodies to distinct epitopes, multiple siRNA’s to knockdown expression, and peptide blocking of nestin immunostaining.

We find that nestin localizes predominantly to the distal axon, and is not found enriched at dendritic tips. In the distal axon, nestin is found in tightly bundled filaments which co-distribute with other neuronal IFs at the wrist region of the growth cone, but individual nestin filaments can also be seen to extend into the central region of the growth cone and even sometimes into the periphery. Nestin is thus positioned to regulate growth cone dynamics.

A remaining question is if the nestin protein detected in neurons is a remnant that was translated when the cell was a NPC, or is made from nestin mRNA in neurons. Our ability to knock down nestin with siRNA following dissociation suggests that nestin mRNA is being actively transcribed after dissociation. Nestin mRNA in neurons would be an indicator that nestin is actually being translated in early neurons. Several paper have described nestin mRNA in neurons in developmental, normal adult, and pathological/injury contexts (Crino & Eberwine 1996; Zhu et al. 2011; Kuo et al. 2005; Arner et al. 2015; Perry et al. 2012; Farzanehfar, Lu, et al. 2017; Dey et al. 2017; Farzanehfar, Horne, et al. 2017; Bigler et al. 2017), thus nestin expression is not excluded from neurons. Nestin’s presence in subpopulations of adult neurons suggests that some neurons may never lose nestin expression, or may regain it later. The best evidence for nestin mRNA in early post-mitotic cortical neurons arises from a single cell mRNA seq experiment, in which flash tag labeling was used to birthdate newborn cells in the cortex of an E14 mouse embryo *in utero*, followed by dissociation and single cell mRNA seq 6 to 48 hours after labeling. This unbiased single cell approach *in vivo*, revealed that nestin mRNA does persist in some newborn neurons of the embryonic mouse cortex (Telley et al. 2016) 24-48 hours after cell division. This suggests that early neurons continue to make new nestin, which we presume still plays a functional role. Furthermore, the initial nestin mRNA localization study did detect nestin mRNA in the intermediate zone (Dahlstrand et al. 1995), where early axons are located.

### A role for neuronal nestin in vivo?

In addition to nestin’s presence in immature neurons, we additionally show that nestin modulates responsiveness to Sema3a. What role could the neuronal nestin play in vivo? At these early developmental time points, Sema3a is important for initial axon positioning and fasciculation (Zhou et al. 2013; Beher er al. 1996). Since Sema3a exists *in vivo* as a gradient emanating from the ventricular zone and cortical plate, it is conceivable that nestin mediated Sema3a sensitivity could dictate the position of the axon within the IZ (Ruediger et al. 2013). A Nestin knockout mouse has been generated previously by others, but was not analyzed for axon guidance defects. The nestin KO mouse is grossly normal except for defects at the neuromuscular junction. Any effects on cortical neuron axon guidance would presumably be subtle, similarly to the Sema3a knockout mouse which is grossly normal with subtle defects in axon fasciculation (Catalano et al. 1998). More detailed analysis of the nestin-positive axons *in vivo* will be needed to more precisely predict what guidance defects might be found if neuronal nestin expression was disrupted. Given our results, this is something of great interest to do in the future, but a conditional KO might be needed to avoid depleting nestin in NPCs, in order to separate any effects of nestin in NPC’s versus in early neurons. We were unable to deplete nestin *in vivo* acutely, because nestin depletion by *in utero* shRNA electroporation, results in loss of migratory neurons in the cortex. Since nestin expression is much higher in NPCs, the authors conclude this phenotype is likely due to impaired cell division of NPCs (Xue & Yuan 2010), precluding analysis of nestin function in postmitotic neurons.

### Nestin as a novel gain control element in Sema3a signaling

Several other molecules have been shown to modulate responsiveness to guidance cues (Kaplan et al. 2014), including types of co-receptors (Dang et al. 2013), lipid raft components (Carcea et al. 2010), second messengers (such as cAMP and calcium) (Henley et al. 2004; Tojima et al. 2014), and modification of downstream signaling cascades by ERM (Mintz et al. 2009), α2-Chimaerin (Brown et al. 2004; Ip et al. 2011), and 14-3-3 proteins (Yam et al. 2012; Kent et al. 2010). Our results now add the IF protein nestin to this category of proteins that can variably regulate responsiveness to Sema3a. The previously identified modulators often serve to vary the degree of signaling output to alter the cytoskeleton and adhesive properties of the growth cone (Vitriol & Zheng 2012). Nestin might similarly modulate cytoskeletal responses to Sema3a, but the molecular mechanism is currently not known. Multiple cytoskeletal proteins are known substrates for cdk5 and act downstream of Sema3a. They are obvious candidates for being nestin effectors in this cascade and mediating the increased Sema3a sensitivity of nestin-expressing neurons. Studies on the role of nestin in cancer cells and other cell types has demonstrated the importance of nestin for migration/invasion, proliferation, and survival-which are reduced after nestin depletion by altering cell signaling pathways (Zhao et al. 2014; Hyder et al. 2014; Narita et al. 2014; Liang et al. 2018; Wei et al. 2007; Zhang et al. 2017; Yan et al. 2016; Matsuda 2013). How nestin promotes higher sensitivity to low doses of Sema3a remains unknown, and future investigation will assess how phosphorylation of cdk5 substrates are affected by the presence of absence of nestin.

One other remaining question is how nestin, which is primarily localized to the distal parts of the axon and to the central domain of the growth cone, affects the stability and dynamics of peripherally localized filopodia. Other cytoskeleton regulating proteins downstream of Sema3a (such as Crmp2, Lis1, tau) (Arimura et al. 2005; Grabham et al. 2007; Astle et al. 2011) also affect filopodia while being primarily localized to the distal axon and the central region of the growth cone. We speculate that microtubules, which can extend in a continuous fashion from the wrist/central region all the way into filopodia, may be directly regulated by wrist-localized effectors. We do not yet know how nestin is affecting filopodia stability downstream of Sema3a, either directly or through other effectors, but this is an exciting future direction for further research.

In summary, we have identified nestin intermediate filament at the axonal growth cone where it plays a role in regulating growth cone morphology. In addition, we identify nestin as a novel downstream effector of Sema3a signaling. Nestin intermediate filament protein could thus play critical regulatory roles in axon guidance responses by serving as a “gain control” element to increase responsiveness of a subset of axons to a common guidance cue.

## Acknowledgements

We thank Brenna Kirk and Pranaya Pakala (Litwa Lab) for work with human IPSCs, Meheret Kinfe and Lloyd McMahon for blinded neuron counting and morphology analysis, Deppman lab for TUJ1 antibody, Laura Digilio, Lloyd McMahon, Kelly Barford (Winckler Lab), and Austin Keeler (Deppman Lab) for help with manuscript revision and expertise, Zdenek Svindrych of the University of Virginia Keck Center for assistance with STED microscopy, and Bruce Lahn’s lab of the University of Chicago for generously providing mouse nestin cDNA. This work was supported by NIH grant R01NS081674 (to BW). CB was supported by an institutional NIH training grant (T32 GM008136).

The authors declare no competing financial interests.

CB and BW conceived and coordinated the study, interpreted the data, and wrote the paper. CJ and KA devised, performed and analyzed Human IPSC studies (Figure 2B and S2A). CB devised, performed, and analyzed all other experiments. CY and ND provided critical expertise for experimental design and execution.

## Material and methods

### Antibodies

Mouse anti-Nestin Rat401-recognizes the unique C-terminal tail region of nestin(Su et al. 2013)–All experiments shown with 2q178 were also repeated with this antibody.

Note this antibody and the other antibodies recognize full length nestin. 1:200 IF, 1:500 IHC, 1:500 WB **DSHB Cat# Rat-401 also rat-401 RRID:AB_2235915**

Mouse anti-Nestin 2Q178 This clone and RAT401 produce indistinguishable staining patterns. All staining and blotting is done with this antibody unless otherwise noted. 1:200 IF, 1:500 IHC, 1:500 WB **Santa Cruz Biotechnology Cat# sc-58813 RRID:AB_784786**

Goat anti-Nestin R-20: Raised against peptide in the unique C-terminal tail region of nestin 1:500 IF, 1:200 IHC **Santa Cruz Biotechnology Cat# sc-21249 RRID:AB_2267112**

Chicken anti-Nestin-three peptide directed antibodies combined to the unique C-terminal tail region of nestin 1:300 IF, 1:400 IHC, 1:3000 WB **Aves Labs Cat# NES RRID:AB_2314882**

Rabbit anti-Nestin AB92391: Raised against peptide in the unique C-terminal tail region of nestin 1:500 IF **Abcam Cat# ab92391, RRID:AB_10561437**

Rabbit anti-DCX-1:1200 IF, IHC, 1:4000 WB **Abcam Cat# ab18723 RRID:AB_732011**

Mouse anti-βIII Tubulin Tuj1-1:800 IF A generous gift from the Deppman Lab YVA **A. Frankfurter, University of Virginia; Department of Biology Cat# TuJ1 (beta III tubulin) RRID:AB_2315517**

Chicken anti-βIII tubulin-1:200 IF **Aves Labs Cat# TUJ RRID:AB_2313564**

Rat anti α-tubulin-1:2000 WB **Santa Cruz Biotechnology Cat# sc-53030 RRID:AB_2272440**

Rabbit anti-Vimentin-1:100 IF, 1:100 IHC, 1:1000 WB **Bioss Inc Cat# bs-0756R RRID:AB_10855343**

Rat anti-Myc tag 9E1-1:1000 WB ChromoTek Cat# 9e1-100, RRID:AB_2631398 Mouse anti-NeurofilamentM 2H3-1:200 IF **DSHB Cat# 2H3 RRID:AB_531793**

Mouse anti-αInternexin 2E3-1:100 IHC **Sigma-Aldrich Cat# I0282 RRID:AB_477086**

Rat anti-L1-1:150 IHC **Millipore Cat# MAB5272 RRID:AB_2133200**

Mouse anti-Tag1-4D7 (DSHB) 1:150 IHC **DSHB Cat# 4D7 RRID:AB_2315433**

Rabbit anti-Calretinin-1:600 IHC **Millipore (chemicon) Cat# PC254L-100UL RRID:AB_564321**

Goat anti-Sox2-1:250 IF **Santa Cruz Biotechnology Cat# sc-17320, RRID:AB_2286684**

Rat anti-CTIP2-1:400 IF **BioLegend Cat# 650601 RRID:AB_10896795**

Rabbit anti-SatB2-1:400 IF **Abcam Cat# ab92446 RRID:AB_10563678**

GFP booster nanobody-1:600 IF **ChromoTek Cat# gba488-100 RRID:AB_2631434**

Phalloidin 568 1:150 Invitrogen

### Neuronal Culture

Primary cultures of cortical neurons were obtained from embryonic day 16 (E16) mouse cortex of either sex as described as approved in the institutional ACUC protocol #3422. Cells were plated on poly-L-lysine coverslips and incubated with DMEM medium with 10% fetal bovine serum. After 8 hrs, the cells were transferred into serum-free medium supplemented with B27 (Invitrogen) and Glutamine and cultured for a total of indicated time periods *in vitro* (DIV). Cells were plated at ~100K cells per 15mm coverslip, and at 50K in the Low Density experiment, or 125k/coverslip in electroporation experiments. Cortical neurons for culture were harvested at E16 prior to glio-genesis-thus non-neuronal glia are rare in these cultures, but occasional nestin/Sox2 positive, GFAP negative persisting neural stem cells were occasionally found.

2d explants were acquired from E16 cortex, but only partially dissociated before plating as described above.

Cells for Sema3a treatment assay were prepared as described above. At 24 or 36 hours, cell were treated with mouse Sema3a (R&D systems) diluted in growth media for 5 minutes at 1nM, or 15 minutes at 2nM as indicated prior to fixation. In some experiments, cells were pre-treated with 10µm Roscovitine (Cayman Chemical) for 30 minutes prior to Sema3a treatment. Time course experiments were fixed at the indicated time points after plating.

### Immunofluorescence

Cells were fixed in 4% Para-PHEM-Sucrose (PPS) for preservation of the cytoskeleton and cellular morphology with prewarmed fixative to 37° C and fixed at 37° C for 20 min. 60mM PIPES, 25mM HEPES, 10mM EGTA, 2mM MgCl2 (PHEM), .12M sucrose, 4% paraformaldehyde pH 7.4. Thorough permeabilization was required for nestin/intermediate filament visualization in neurons. Coverslips were permeabilized with 0.4% triton-X 200 in 1% BSA PBS for 20 min for nestin and intermediate filament staining, or 0.2% triton-X when no intermediate filaments were being immunostained, and then blocked for 30 min in 10%BSA, 5% normal donkey serum (NDS) in PBS. Primary antibodies were diluted in 1% BSA in PBS, and secondary antibodies in 1%BSA, 0.5% NDS in PBS. Primary antibody were incubated overnight at 4° C, and secondary antibodies for 90 min at room temperature. Appropriate species specific alexa 350, 488, 568, or 647 labeled secondary antibodies raised in Donkey (Invitrogen) were used for fluorescent labeling. Phalloidin 568 (Invitrogen) at 1:150 was used to visualize F-actin and DAPI to visualize Nuclei. For peptide blocking negative control experiments, the goat anti-nestin antibody was diluted at 1:500 with the corresponding epitope immunizing peptide (Santa Cruz) at 1:50 and incubated at room temperature for 3 hours. The goat anti-nestin antibody for the non-blocked condition was diluted and incubated similarly but in the absence of peptide. Coverslips were mounted with ProLong Gold(Invitrogen) and imaged with a Zeiss Z1-Observer with a 40x (for population counting) or 63x (for morphology analysis) objective. Images were captured with the Axiocam 503 camera using Zen software (Zeiss) and processed identically within experiments. No non-linear image adjustments were performed.

### Histology

Human IPSC derived Cortical Organoids (minibrains) or embryonic E16 mouse brain were dissected and placed in 4% PFA and 6% sucrose in PBS at 4C overnight. Tissue was then cryo-protected by sinking overnight in 30% sucrose. Tissue was imbedded in OCT, and brains cut by coronal cryosection at 16um sections (Zeiss Confocal imaging) or 10um sections (Abberior super-resolution). Sections were mounted directly on slide, permeabilized with 3% NDS + 0.2% triton-X 100 in PBS for 1 hour, and blocked with 10% BSA for 30 min. Antibodies were diluted in permeabilization buffer with indicated primary antibodies overnight at 4° C, and secondary antibodies 2 hours at room temperature following thorough washing with PBS. Sections were mounted with prolong gold. For peptide blocking negative control experiments, the goat anti-nestin antibody was diluted at 1:100 with the corresponding epitope immunizing peptide (Santa Cruz) at 1:10 and incubated at room temperature for 3 hours. The goat anti-nestin antibody for the non-blocked condition was diluted and incubated similarly but in the absence of peptide. The antibody was then used as described above. Of note, the anti Tag1 IgM antibody has cross reaction with mouse IgG secondary antibodies, so staining with mouse IgG and mouse IgM antibodies were done sequentially. Confocal imaging of cryosections was carried out on an inverted Zeiss LSM880 confocal microscope using a 63X objective. Super-resolution STED imaging utilized and Abberior STED confocal with 100X objective at the Keck Center and University of Virginia. A donkey ani-goat 488 (Invitrogen) secondary antibody was imaged using a 595 nm extinction laser.

### Neuron nucleofection

After cell were dissociated from cortex, but prior to plating, cells were electroporated by Amaxa 4d nucleofection, according to protocol. 800,000 cells were electroporated in the small volume (20µl) cuvettes using P3 primary cell solution and EM-100, CL-133, and CU-100 programs variably between experiments. All three of these programs worked similarly in our hands. 0.1µg of GFP plasmid (pCMV-GFP-Clonetech) as a transfection marker, and 100nM siRNA (siCON-SCBT sc-37007, siNes smart-pool Dharmacon siGenome M-057300-01-0010, consisting of 4 individual siRNA’s (D-057300 -01,-03,-04, and -17) with target sequences: siRNA #1-AAAGGUGGAUCCAGGUUA, siRNA #3 D-AGACAAGACUCAUGAGUC, siRNA #4 CAGCAGGCAUUUAGACUUC, siRNA #17 GCGACAACCUUGCCGAAGA). All 4 target the region in the mRNA corresponding the unique c-terminal tail of nestin protein, which is unlike other IF’s.

### Molecular biology

Circular mouse nestin cDNA was obtained as a gift from Bruce Lahn. Briefly, PCR was used to generate ECOR1 and HINDII sites on the 5’ or 3” end respectively of full length mouse nestin. Both the insert and vector were treated with EcoR1 and HindII, the cut insert was ligated into pCDNA3.1 Myc-His(+) B.

### Cell culture and transfection

HEK293 cells were maintained in DMEM+10% fetal bovine serum, and all transfections were conducted using Lipofectamine 2000 (Invitrogen) according to the manufacturer's protocol. mNestin-myc and respective control or nestin siRNAs were transfected simultaneously according to the recommended instructions.

### Western Blot

Cells were washed with ice cold PBS, and lysed directly on the plate (6cm) into lysis buffer(400ull); 50mM Tris HCl, 150mM NaCl, 1% NP-40, 1mM EDTA, 1mM DTT, 1mM PMSF, 1X HALT protease/phosphatase inhibitor cocktail. After rotisserie at 4° C for 30 minutes, lysate were cleared by centrifugation at 21K X g for 15 min at 4° C. Supernatants were diluted with 5X laemmli buffer and equivalent amounts were run on a 4-20% polyacrylamide gel. After transfer onto nitrocellulose with the BioRad Transblot, membrane was blocked with Licor TBS blocking buffer for 1 hours at room temp. Primaries and secondary antibodies were diluted in 50% blocking buffer, 50% PBS, 0.1% tween-20. Primaries overnight incubation at 4° C, and near-infrared secondary antibodies 1 hour at room temperature. Blot were imaged with the Licor Odyssey CLx imager.

### Image analysis

Image J was used for all image analysis

*Neuron morphology and nestin expression analysis*. Neuron staging was performed according to the Banker neuron staging method, and neurites were called an axon if it is 2X longer than the next longest neurite (dendrite). Nestin positivity in the axon of neurons was assessed by outlining the distal region of the axon and measuring the maximum nestin immuno-staining in the region. If this value was at least 1.5x above average background, then a cell would be considered positive for nestin. In siRNA experiments, only green GFP+ (transfection indicator) cells were imaged and counted for both for morphology and nestin siRNA efficiency analysis. All counts were confirmed by blinded analysis. Depiction of how morphology measurements were made is in Figure S4F.

Filopodial protrusion were counted using the phalloidin channel, and any actin rich protrusion from the growth cone was counted.

Axon length was the length of the axon from the cell body to the growth cone tip. Growth cone area and perimeter were assessed by measuring traced outlines the growth cone not counting the filopodial protrusions, and extending down to the point where the splayed growth cone microtubules consolidate into bundles in axon shaft. Mouse Neural stem cell marker expression. DCX, nestin, and DAPI stained differentiated neural stem cells were manually counted for immunopositivity in any part of the cell above background levels.

Human IPSC Nestin and βIII tubulin co-localization was assessed per image using the ImageJ co-localization plugin.

### Mouse neural stem cell culture

Embryonic (E16) cortical neural stem cells were prepared as described(Pacey et al. 2006) and grown as neurospheres for at least 3 passages prior to use for dissociated and differentiating cell experiments to ensure purity. For these experiments, neurospheres were dissociated using Stem Cell chemical dissociated kit, and plated in PLL coated coverslips for 1 day and fixed and immunostained as described. For 4DIV, proliferative media was replaced with media lacking EGF and FGF growth factors until 4 DIV, fixed and processed for immunostaining as described.

### Neural Differentiation of Human Induced Pluripotent Stem Cells

Human induced pluripotent stem cells from neurotypic control 9319 were generated by Kristen Brennand in the laboratory of Dr. Fred Gage (Salk Institute). Neural progenitor cells (NPCs) were generated through an embryoid body protocol as described by Brennand et al. PMID: 21490598. For neuronal differentiation, we plated 75,000 control NPCs on each poly-ornithine and laminin-coated 18mm glass coverslip. After 24 hours, cells were switched to neuronal differentiation media consisting of DMEM F12 + Glutamax and supplemented with N2 and B27 with retinoid acid and 20ng/ml BDNF, 20ng/ml GDNF, 400uM cyclic AMP, and 200nM ascorbic acid. Half of the media was replaced with fresh media every 3 days. Neurons were fixed with 4% formaldehyde with 4% sucrose on the days indicated in the text and used for subsequent immunofluorescence analysis.

### Generation of Cortical Organoids

Human fibroblasts from neurotypic control 7545 were obtained from the Coriell Institute and reprogrammed into induced pluripotent stem cells by the laboratory of Dr. Mike McConnell (UVa) using the CytoTune Reprogramming Kit (Life Technologies) and maintained in Essential 8 media (Life Technologies). Cortical organoids were grown using a low-attachment protocol^20^. Briefly, we used dispase to detach hIPSC colonies from matrigel dishes. These colonies were placed in ES/DMEM media (GlobalStem) supplemented with 20% Knockout Serum Replacement (Life Technologies) and treated with 5mM Dorsomorphin and 10mM SB-431542 for the first 5 days. The ROCK inhibitor, 10mm Y-27632, was added for the first 48 hours. From day 6-24, spheroids were fed with Neuronal Media consisting of Neurobasal A with B27 without Vitamin A, GlutaMax, penicillin/streptomycin (Life Technologies), and supplemented with 20ng/ml EGF and FGF2 (Peprotech). From day 25-42, the growth factors were replaced with 20ng/ml BDNF and NT3 (Peprotech). From Day 43 onwards, BDNF and NT3 were removed and organoids were grown solely in Neuronal Media.

### Statistical analysis

Human IPSC Nestin and β3 tubulin colocalization was performed using Sigma Plot 13.0. All other statistical analysis was carried out using Prism software. The datasets were first evaluated as parametric versus nonparametric using the Shapiro-Wilk normality test. The corresponding one way ANOVA test was used when there were multiple comparisons, and the t-test was used when only two conditions were compared, also as indicated in figure legends.

