## Supplementary material for "Nestin in immature embryonic neurons regulates axon growth cone morphology and Semaphorin3a sensitivity": Online Supplemental Material

FigureS1A and A' demonstrate the knockdown efficiency of nestin specific siRNAs used in this study on endogenous nestin in E16 cortical neurons (A), and on overexpressed myc- tagged nestin in Hek293 cells (A'). FigureS1B and B' show full length western blots probed with both the mouse anti-nestin and goat anti-nestin antibodies used in this study. Lysates from E16 1DIV neurons in culture (B), and lysates of E16 whole cortex (B') were used. FigureS1C demonstrates the relative expression level of neuronal nestin in DCX positive stage3 E16 mouse neuron compared to the high nestin expression in a Sox2 positive neural stem cell.

FigureS2A demonstrates nestin expression in βIII-tubulin positive neurons in a human IPSC derived cortical minibrain. FigureS2B shows that both Ctip2 and Satb2 neurons have nestin expressing populations.

FigureS3A demonstrate additional nestin immunostaining of mouse E16 cortical coronal cyrosection with both the mouse anti nestin 2Q178 clone, and the classic mouse anti-nestin Rat401 clone. The staining was carried out as quadruple immunostainings in which the mouse anti-nestin antibody staining was combined with another nestin antibody (i.e. goat anti-nestin), antibody against vimentin (a polymerization partner for nestin) antibody, and antibody against the axonal adhesion molecule L1-CAM.

FigureS4 serves as validation for the nestin siRNA's used in this study. Figure S4A determines the effectiveness of each of the 4 nestin siRNA's, both separately and as a pool, at depleting over-expressed myc- tagged nestin protein in Hek293 cells. siRNAs 1 and 17 were equally effective as the siRNA pool to deplete nestin in neurons (FigureS4B). siRNAs 1 and 17 have no effect on axon length (figureS4C), but cause an increase in growth cone area (FigureS4D), and an increase in growth cone filopodia number (FigureS4E). FigureS4F illustrates how growth cone area, growth cone filopodia number, and axon length were measured.

**Supplementary Figure 1**

A. Nestin knockdown was also qualitatively assessed by western blotting of whole cell lysates of cortical cultures (1 DIV) with Mouse, Goat, and Chicken nestin antibodies. A band of the expected size was detected with all 3 antibodies in siCon but decreased levels are seen after siNes expression. The knockdown was partial. This is likely due to lower transfection efficiencies in cultured neurons where less than 50% of cells are transfected. The related intermediate filament vimentin was unaffected, and DCX was blotted as a loading control.

A'. Myc-tagged nestin was overexpressed in HEK293 cells and co-transfected using lipofectamine2000 with siCon or siNes siRNA. Lysates were blotted for the myc-tag and for nestin, demonstrating near complete knockdown in the high transfection efficiency HEK293 cells. The related intermediate filament vimentin was unaffected, and alpha-tubulin is loading control.

B and B'. Full blots demonstrating specificity of nestin antibodies used in IF studies as a single band above 250 kd. Protein lysate from cultured 1DIV E16 mouse neurons (B), and E16 mouse cortex (B'), both probed with the goat anti-Nestin antibody and the mouse anti-Nestin clone 2q178 antibody.

C. Side by side comparison of nestin levels in a Sox2 positive neural progenitor cell (NPC) and a primary differentiating DCX positive neuron. E16 mouse cortical cultures at 1DIV were stained with antibodies to the indicated proteins. They contain mostly neurons but an occasional NPC can still be found at 1 DIV. Nestin levels in the in the primary neurite and cell body of DCX+ neurons are many fold lower than in the NPC, but is still detectable.

**Supplementary Figure 2**

A. Nestin is expressed in βIII tubulin-positive neurons in human IPSC derived mini-brains. The boxed regions 1-3 are shown larger in the insets, and arrowheads indicate nestin βIII tubulin doublepositive cells. Nestin positive radial glia like cells span the spheroid, while high βIII tubulin cells are located to the periphery. Low βIII tubulin expressing cells can also be found in the central ventricular-like region representing differentiating neurons not yet migrated, and insets analyzed are outside of this region. Nestin can be detected in the tips of some of the βIII tubulin processes (arrowheads).

B. Both nestin negative and nestin positive have a range of Ctip2 and Satb2 nuclear intensities. Pink arrowheads indicate nestin in the distal axon. A total of 68 DCX+ neurons (24 nestin negative, 44 nestin positive) were quantified in terms of nuclear intensity of either Ctip2 or Stab2 immunostaining.  No correlation was found. (Unpaired t-test)

**Supplementary Figure 3**

A,B.  Nestin (multiple antibodies) is found together with its polymerization partner vimentin along axons (identified by the axonal cell adhesion molecule L1-CAM). Arrow heads indicate radial glia and arrows indicate nestin-positive axons.
Both the mouse anti-nestin antibody 2q178 (A) and rat401 (B) produce similar axon immunostaining in E16 mouse cortex, as do the other nestin antibodies used in this study (Figure 3b).

**Supplementary Figure 4**

A.  Relative efficiency of nestin silencing of the 4 individual siRNA's from the siNes pool. Myc-tagged nestin was overexpressed in HEK 293 cells and co-transfected using lipofectamine 2000 with siCon or various siNes siRNA.  Lane 1 is untransfected HEK293 cell lysate.  Lysates were blotted for the myc-tag and for nestin, demonstrating again complete knockdown with the siNes pool, as well as efficient knockdown by siNes #1 and 17.  siNes #3 had an intermediate effect, while siNes #4 was the least effective, and is used as an additional control in the following experiments in neurons.    The related intermediate filament vimentin was unaffected, and □-tubulin is used as a loading control.

B-E. 3 individual siNes siRNA's were transfected into neurons to confirm efficiency (B) and confirm morphological phenotypes (C,D,E) seen after nestin depletion by siNes pool.  Both effective siNes #1 and #17 (and the siNes-pool) resulted in a significant depletion of nestin positive neurons after 36 hours, while the non-effective siNes #4 did not significantly reduce the number of nestin positive cells.  None of the transfection conditions altered the average axon length (C), while siNes #1 and #17 phenocopied the pool by growth cone area (D) and growth cone filopodia number (E), while siNes #4 had no significant effect.  Error bars are SEM.  N=3 experiments, with the means of each plotted on the graph. Normality was confirmed with the shapiro-Wilk normality test. Each condition was compared to control with a one way ANOVA with Dunnet's correction.

F. A labeled version of the cell in Figure 5E diagraming how morphological measurements were made.
