## Supplementary figures and images for "Nestin in immature embryonic neurons regulates axon growth cone morphology and Semaphorin3a sensitivity"

### SupplementFig1

#### Supplementary Figure 1

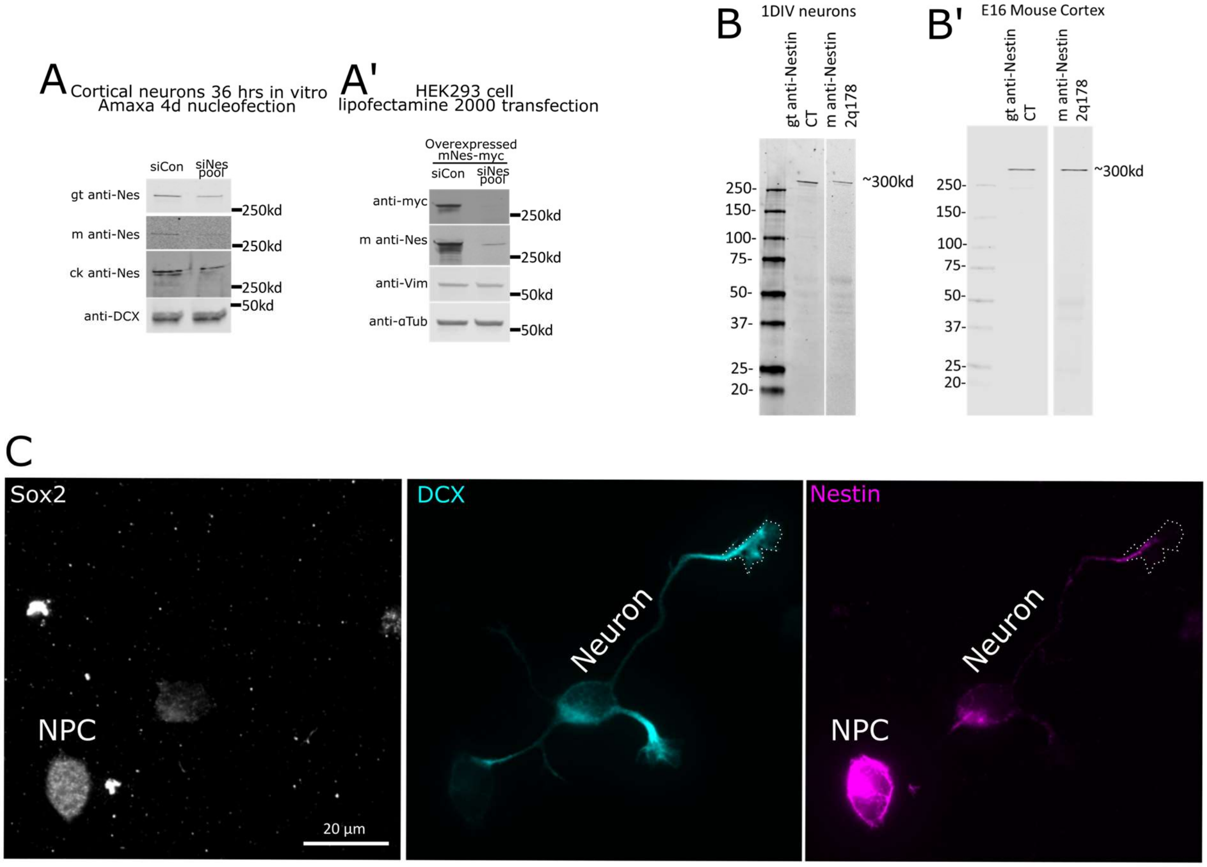

### SupplementFig2

#### Supplementary Figure 2

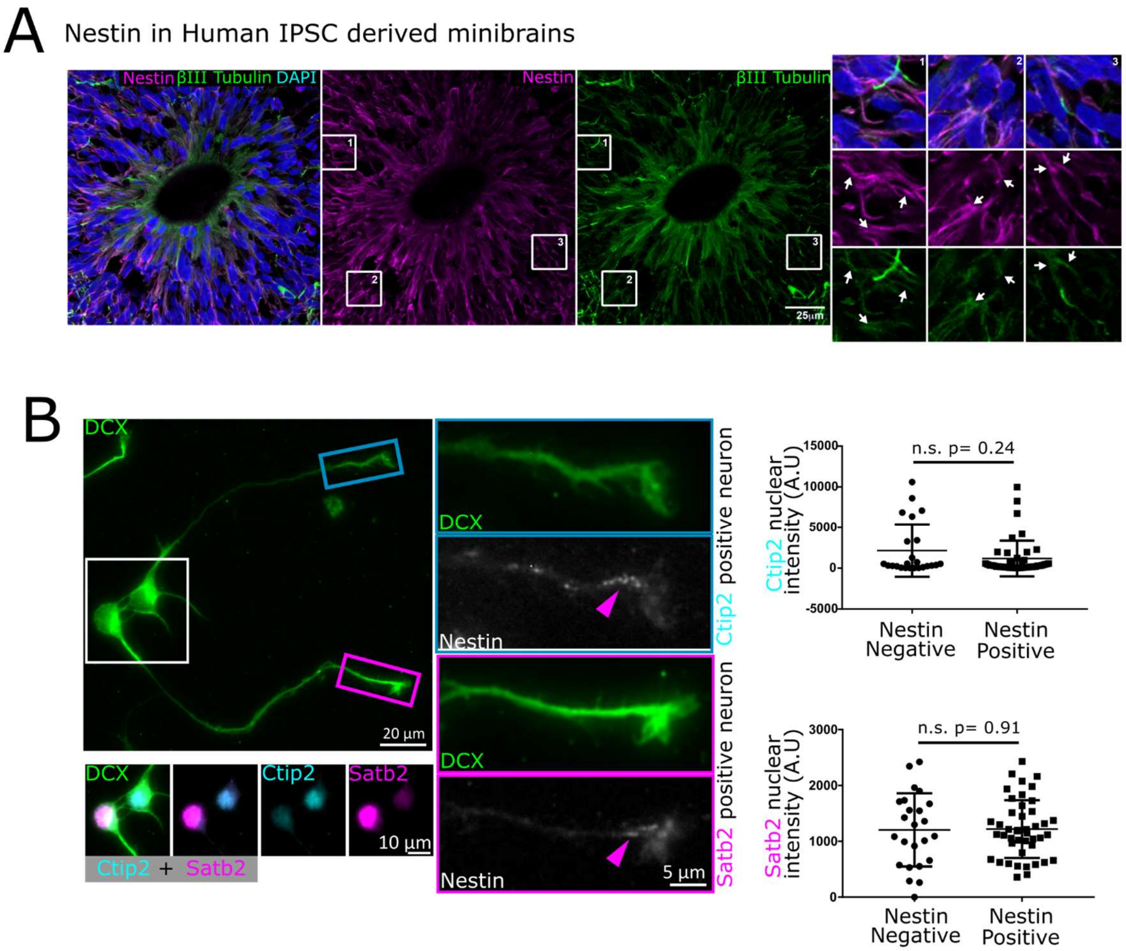

### SupplementFig3

#### Supplementary Figure 3

A

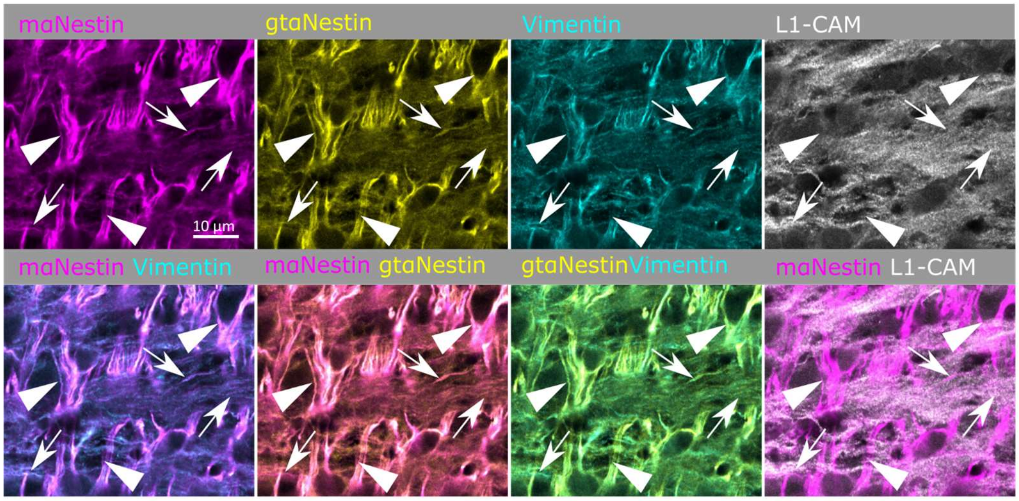


B

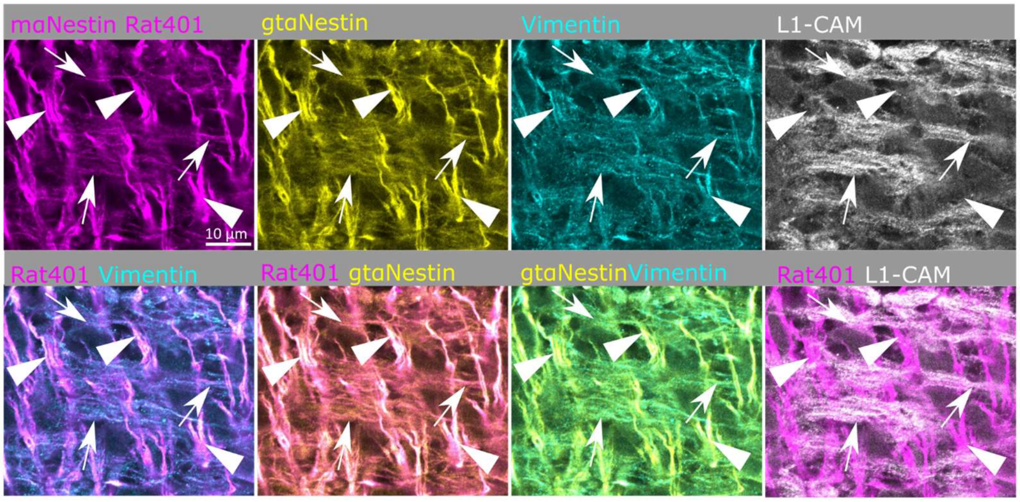

### SupplementFig4

#### Supplementary Figure 4

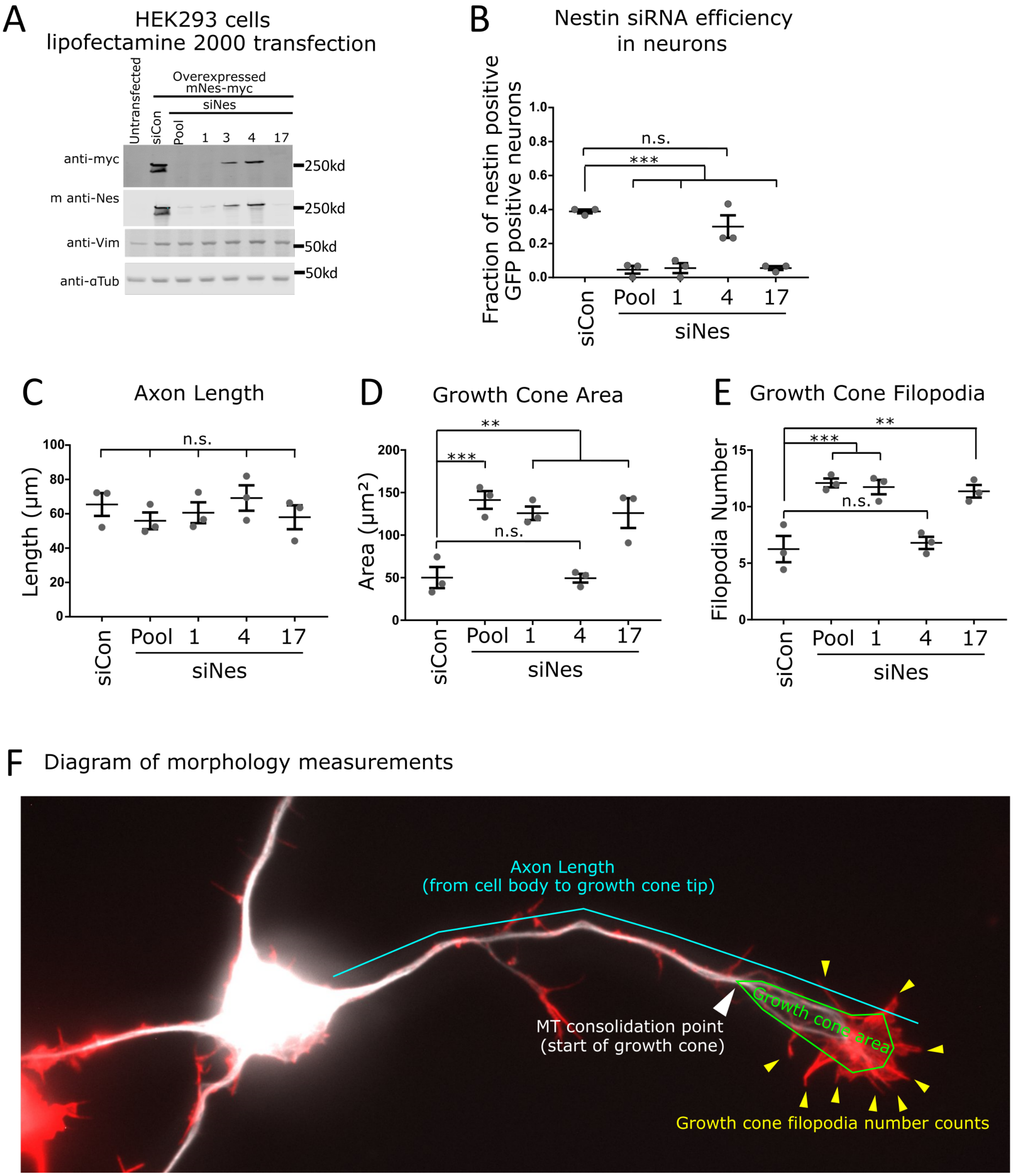
